# *Streptococcus pyogenes* M1T1 variants activate caspase-1 and induce an inflammatory neutrophil phenotype

**DOI:** 10.1101/2020.03.02.972240

**Authors:** Jonathan G. Williams, Diane Ly, Nicholas J. Geraghty, Jason D. McArthur, Heema K. N. Vyas, Jody Gorman, James A. Tsatsaronis, Ronald Sluyter, Martina L. Sanderson-Smith

## Abstract

Invasive infections due to Group A *Streptococcus* (GAS) advance rapidly causing tissue degradation and unregulated inflammation. Neutrophils are the primary immune cells that respond to GAS. The neutrophil response to GAS was characterised in response to two M1T1 isolates; 5448 and animal passaged variant 5448AP. Neutrophil co-incubation with 5448AP allowed proliferation of GAS while it also lowered the production of reactive oxygen species by neutrophils when compared with 5448. Infection with both strains invoked neutrophil death, however apoptosis was reduced in response to 5448AP. Both strains induced neutrophil caspase-1 activation and caspase-4 expression *in vitro*, with caspase-1 activation detected *in vivo*. Thus, GAS infection of neutrophils corresponds to increased caspase-1 activity and caspase-4 expression, consistent with inflammasome activation and pyroptosis. GAS infections that promote an inflammatory neutrophil phenotype may contribute to increased inflammation yet ineffective bacterial eradication, contributing to the speed and severity of invasive GAS infections.

## Introduction

Invasive infections due to the obligate human pathogen *Streptococcus pyogenes* (Group A *Streptococcus*; GAS) are characterised by unregulated inflammation and high mortality (Davies et al., 1996, Lamagni et al., 2008, Nelson et al., 2016, O’Grady et al., 2007, O’Loughlin et al., 2007, Svensson et al., 2000). The highly virulent GAS M1T1 clone 5448 is well studied (Aziz and Kotb, 2008, Tart et al., 2007, Walker et al., 2007). The propensity of M1T1 GAS to cause invasive infections is due, in part, to the increased frequency of spontaneous mutations in the two-component control of virulence regulatory system (covRS) (Walker et al., 2007), resulting in differential expression of multiple virulence factors, many of which are implicated in immune defence (Kilsgård et al., 2016, Sumby et al., 2005, Sumby et al., 2006, Walker et al., 2007). Compelling evidence describes the resistance of *covS* mutant GAS (mutation to the sensor protein of covRS) to killing by human neutrophils (polymorphonuclear leukocytes, PMNs) (Ato et al., 2008, Maamary et al., 2010, Walker et al., 2007). 5448AP (animal passaged) is one M1T1 *covS* mutant strain that shows resistance to killing by human neutrophils, has increased bacterial dissemination and results in decreased survival in mouse models of infection (Fiebig et al., 2015, Maamary et al., 2010, Walker et al., 2007). Due to these characteristics, 5448AP can be used as a model strain to explore the host response to invasive GAS infection.

Neutrophils are the primary innate immune cell-type that defend against bacterial pathogens, with their presence contributing to the overarching inflammatory response (Kobayashi et al., 2018). Proteins of neutrophil origin are highly abundant at sites of invasive GAS infection (Edwards et al., 2018). The neutrophil life cycle is tightly regulated to avoid the release of cytotoxic contents that can damage surrounding tissue and contribute further to inflammation (Epstein and Weiss, 1989, Faurschou and Borregaard, 2003). Under normal conditions cytotoxic reactive oxygen species (ROS) produced by neutrophils kill phagocytosed bacteria (Nguyen et al., 2017). An increased production of ROS signals for the induction of the anti-inflammatory cell-death pathway apoptosis (Lundqvist-Gustafsson and Bengtsson, 1999). During apoptosis, neutrophils are carefully decommissioned, ensuring no unregulated release of intracellular material, then cleared by a secondary phagocyte, macrophages, through a process known as efferocytosis (Bratton and Henson, 2011). The cleavage of caspase-3 to the p17 form denotes the execution of apoptosis (Nicholson et al., 1995). Increased neutrophil apoptosis is described during GAS infection (Kobayashi et al., 2003). Pro-inflammatory neutrophil death has also been reported during GAS infection (Tsatsaronis et al., 2015). One form of pro-inflammatory death is pyroptosis, involving the formation of an inflammasome and subsequent caspase-1 or caspase-4 activation, culminating in cell-lysis (Man and Kanneganti, 2016, Sollberger et al., 2012). Deregulation of (or excessive) pro-inflammatory neutrophil-death may therefore contribute further to inflammatory pathology, as demonstrated during infection with other pathogens, including *Pseudomonas aeruginosa* (Ryu et al., 2017) and *Staphylococcus aureus* (Ghimire et al., 2018, Greenlee-Wacker et al., 2017, Ryu et al., 2017). Inflammation is also heavily influenced by cell cytokine production, where increased release can invoke neutrophil dysfunction, affecting neutrophil survival, antibacterial function or even migration (Brown et al., 2006, Kipnis, 2013, Kovach and Standiford, 2012, Norrby-Teglund et al., 2000). Previous work outlines how GAS take advantage of inadequate or exacerbated immune responses and focus upon the bacterial mechanisms involved (Cole et al., 2011). Although models to investigate leukocyte inflammasome activation have been proposed (Tran et al., 2019), a limited number of studies have investigated the neutrophil response to GAS specifically (Kobayashi et al., 2003). Here, using an *in vitro* model, we temporally describe the effect two M1T1 GAS strains have upon human neutrophil death, signalling and inflammatory profile. We further characterise this interaction in murine neutrophils, using an *in vivo* model of intradermal invasive GAS infection. Additionally, we identify differences in the neutrophil response to GAS due to *covS* mutation. We hypothesise that changes to neutrophil function by GAS can promote invasive GAS infection by exacerbating the inflammatory response, decreasing antibacterial capacity and inducing inflammatory neutrophil death.

## Results

### GAS persistence and proliferation occurs during a dampened neutrophil ROS response

Neutrophil phagocytic ability is a predominant and well-established mechanism of immune defence. Both GAS strains 5448 and 5448AP rapidly associated with human neutrophils, with near maximal association observed within 30 min (Figures 1A and S1A-C). Higher percentages of neutrophils were associated with 5448 compared to 5448AP (Figure 1A). 5448 and 5448AP also induced ROS production in neutrophils, with less ROS produced by neutrophils incubated in the presence and absence of 5448AP compared to those incubated with 5448 (Figure 1B). As such, the rate of neutrophil ROS production was significantly increased compared to the neutrophil only control during 5448, but not 5448AP, infection (p<0.01, Figure S1D), suggesting dampened antibacterial function in neutrophils in response to 5448AP. This difference in GAS-induced ROS production inversely corresponded to GAS proliferation in the presence of neutrophils, with high numbers of 5448AP observed compared to 5448 over 180 min (Figure 1C). Differential GAS proliferation required viable neutrophils, as 5448 and 5448AP proliferation in the presence of neutrophil lysates was reduced and similar for both strains over 180 min (Figure 1D). Finally, incubation of neutrophils with the phagocytosis inhibitor cytochalasin D significantly increased 5448, but not 5448AP, survival (p<0.001, Figure S1E), indicating that 5448 killing is predominantly mediated via a phagocytic-related pathway. Collectively, these data indicate that compared to 5448, 5448AP associates less readily with human neutrophils leading to reduced ROS production and increased survival of this strain.

**Figure 1.**
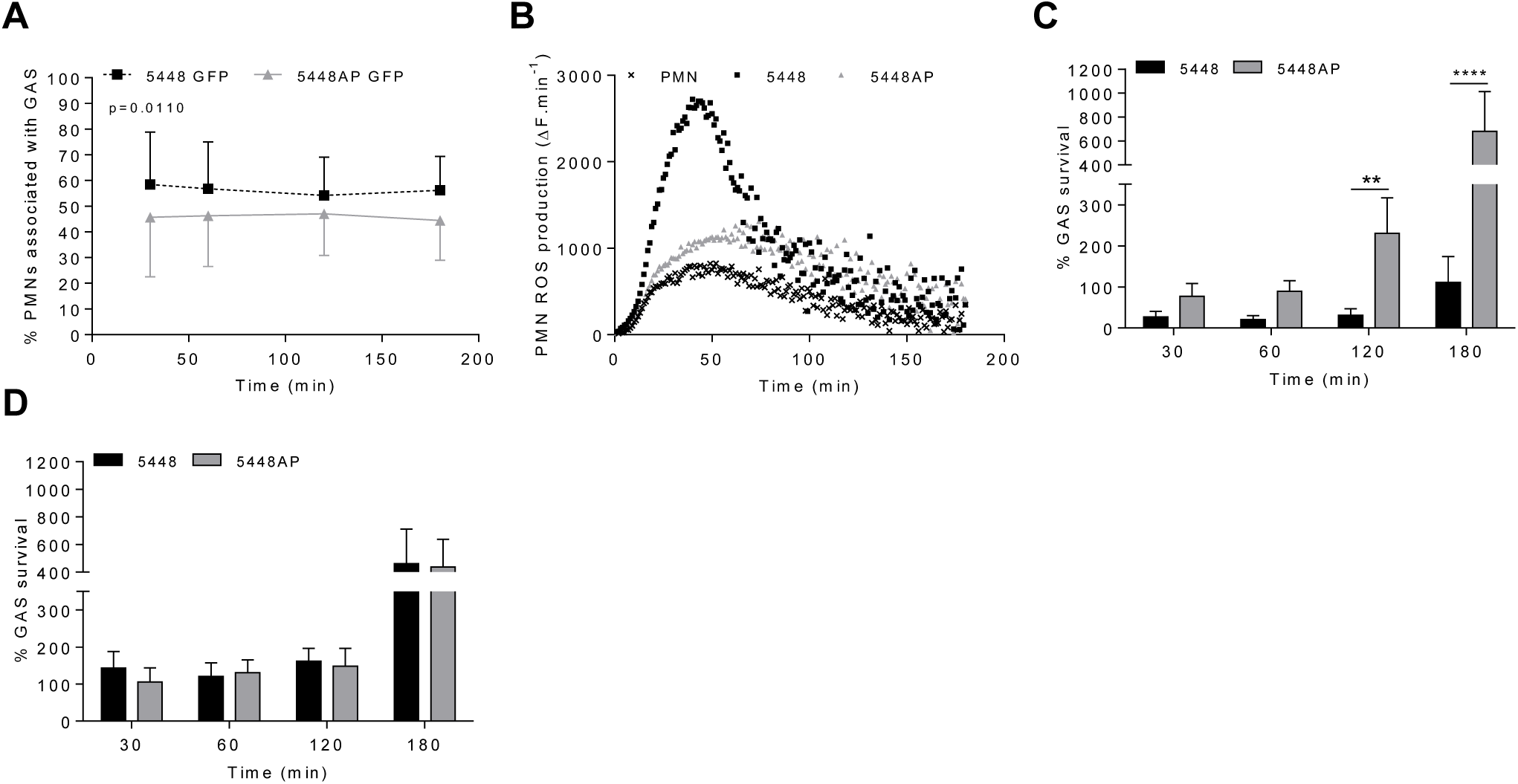
GAS persistence and proliferation occurs during a dampened neutrophil ROS response. (A) Association of fluorescent GAS strains with human neutrophils (PMNs, n=6 donors), analysed via flow cytometry (Figure S1A-B). (B) Infection of human neutrophils with GAS invokes ROS production. Representative of triplicate measurements from three separate experiments (n=3 donors), shown as mean change in fluorescence units over time (ΔF.min^-1^). GAS strains were incubated in the presence of (C) human neutrophils (n=8 donors) or (D) lysed human neutrophils (n=4 donors) over 180 min with surviving bacterial concentration determined as percentage of inoculum. Results are the pooled means±SD (of triplicate measurements for panels C and D). **p<0.01 and ****p<0.0001.

### Neutrophil death is delayed during infection with 5448AP

GAS can induce death of human neutrophils (Kobayashi et al., 2003, Tsatsaronis et al., 2015), however the mechanism remains poorly described. Phosphatidylserine (PS) exposure is a hallmark of the induction of multiple death pathways including apoptosis and pyroptosis (Man and Kanneganti, 2016, Wang et al., 2013). Human neutrophils were incubated with 5448, 5448AP or in the absence of GAS then the binding of annexin-V (AV), as a measure of PS exposure, and the binding of Zombie Fixable Viability Kit (Z), as a measure of membrane integrity, were analysed by flow cytometry (Figure 2). Cells were defined as viable (AV^-^Z^-^), PS exposed (AV^+^Z^-^) or dead (AV^+^Z^+^) (Figure 2C). Neutrophils incubated without GAS remained viable over 180 min (Figure 2D), with 13% exposing PS (Figure 2E) and 3% dead after this time (Figure 2F). In contrast, incubation of neutrophils with either 5448 or 5448AP resulted in increased PS exposure and cell-death over 180 min, with significantly more cell-death during 5448 infection compared to 5448AP infection at 30 and 60 min (p<0.05, Figure 2F). Delay in the initial induction of death in response to 5448AP suggests extended neutrophil viability during the early stages of infection (30-60 min).

**Figure 2.**
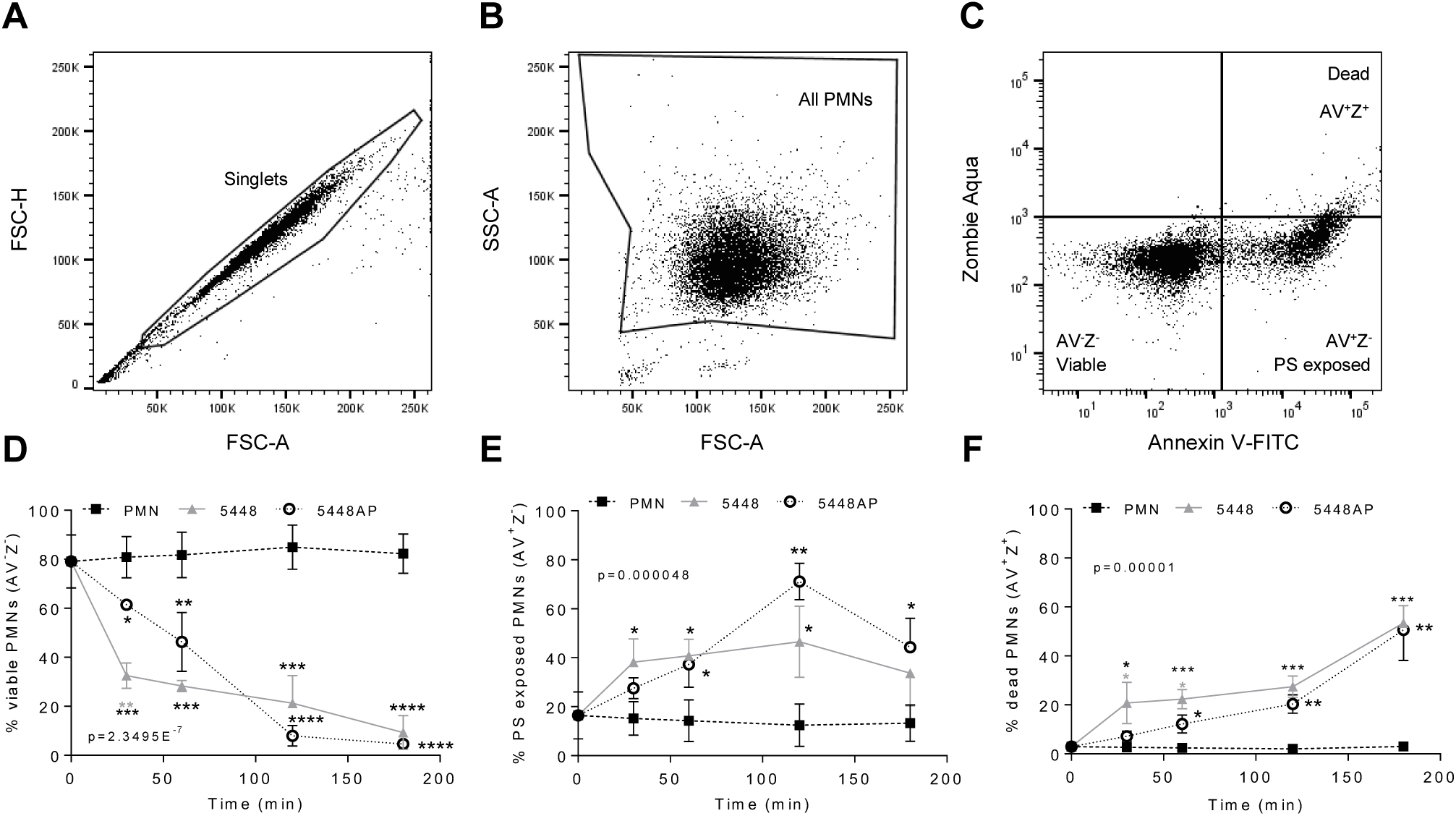
GAS infection induced neutrophil death. Human neutrophils (PMNs) were stained with Annexin V-FITC and Zombie Aqua Fixable Viability Kit via flow cytometry and initially gated upon (A) ‘Singlets’ (FSC-A vs. FSC-H, 10,000 events collected). ‘All PMNs’ (live and dead) were selected using (B) a gate excluding GAS. Cells were then compared for fluorescence of (C) annexin V-FITC/phosphatidylserine (PS) binding against Zombie live/dead viability dye. Purified PMNs were sampled over 180 min to measure (D) ‘Viable’ (AV^-^Z^-^), (E) ‘PS exposed’ (AV^+^Z^-^), and (F) ‘Dead’ (AV^+^Z^+^), neutrophils during GAS infection (n=3 donors). Results are pooled means±SD. *p<0.05, **p<0.01, ***p<0.001 and ****p<0.0001, with black asterisks denoting significance from control and grey asterisks between 5448 and 5448AP.

### GAS infection activates caspase-1 in neutrophils

The above data (Figure 2) suggests that GAS induce rapid neutrophil death. To explore the mechanism of GAS-induced neutrophil death human neutrophil lysates were separated by SDS-PAGE and screened for the abundance of active caspase-3, caspase-1 and caspase-4 via immunoblotting and quantified using densitometry (Figures 3A and S2A-D). Cleavage of executioner caspase-3 to p17 is a definitive hallmark of apoptosis induction (Nicholson et al., 1995). Conversely, caspase-1 and caspase-4 activity can indicate inflammasome activation, which in turn can cleave downstream effectors of pyroptosis (Man and Kanneganti, 2016, Sollberger et al., 2012). Moreover, a recent report has identified the caspase-1 p46 molecule as the principal active species during inflammasome activation in the cell (Boucher et al., 2018). Neutrophils incubated in the absence of GAS showed limited evidence of caspase-3 cleavage to p17 (Figures 3A and S2A), while levels of caspase-1 reduced over 180 min (Figures 3A and S2B-C). Infection of neutrophils with 5448 significantly increased the presence of caspase-3 p17 at 180 min (p>0.0001) compared to both 5448AP-infected and uninfected neutrophils (Figures 3A and S2A). In contrast, incubation of neutrophils with either GAS strain caused an upregulation of caspase-1 (Figures 3A and S2B-C). Compared to uninfected neutrophils, 5448 infection significantly increased pro-caspase-1 (50kDa) at 30 min, 60 min (p<0.05) and 180 min (p<0.01), whilst 5448AP infection only significantly increased pro-caspase-1 at 180 min (p<0.001, Figures 3A and S2B). Further, 5448 and 5448AP infected neutrophils sustain p46 expression over 180 min, though this difference was not statistically significant (Figure 3A and S2C). Significant differences were not seen in the expression of caspase-1 between 5448 and 5448AP infected neutrophils. Similar to caspase-1, the abundance of full-length caspase-4 in neutrophils incubated in the absence of GAS declined over time (Figures 3A and S2D). In contrast, 5448 infection increased caspase-4 expression at 30 (p<0.01), 60 (p<0.001) and 180 min compared to uninfected neutrophils (p<0.0001, Figures 2A and S2D). 5448AP infection however, only increased caspase-4 expression at 180 min (p<0.01, Figures 3A and S2D).

**Figure 3.**
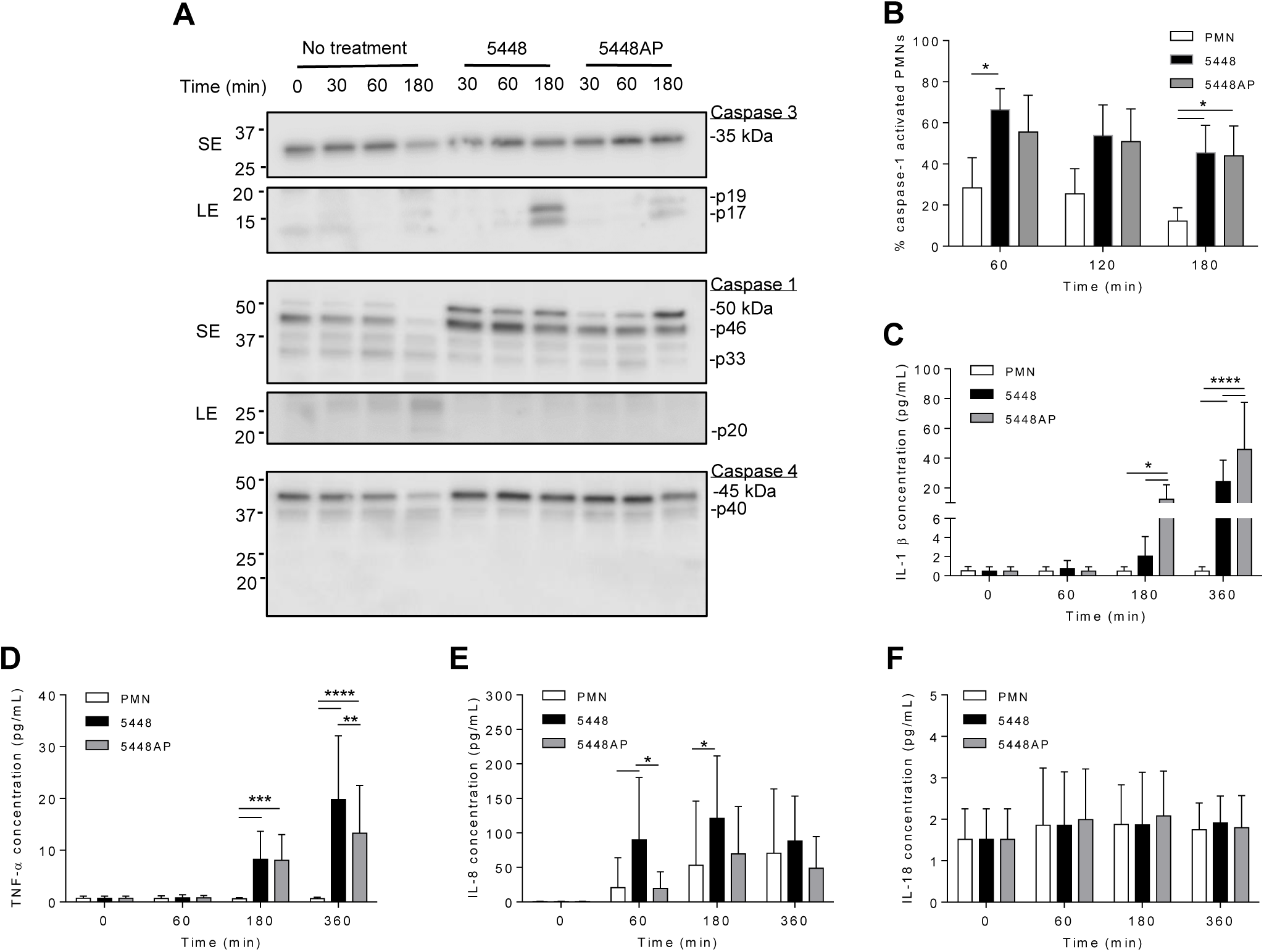
GAS infection activates caspase-1 in human neutrophils and induces proinflammatory cytokine IL-1β and TNF-α release. (A) Human neutrophil (PMN) lysates were prepared at 30, 60 and 180 min during GAS infection and compared to uninfected neutrophils (0, 30, 60, 180 min) via immunoblotting of caspase-3, caspase-1 and caspase-4. Images shown are from a single donor and are representative of triplicate experiments using different donors. Immunoblot bands were quantified (ImageJ) and normalised over total protein (Figure S2). SE=short exposure, LE=long exposure. (B) Caspase-1 activation in human neutrophils due to GAS infection was confirmed via flow cytometry using FLICA (FAM-YVAD-FMK) (n=3 donors). Flow cytometry gating strategy used singlets/PMNs (figure S3A-B) and FAM-FLICA fluorescence (Figure S3C). The release of cytokines from neutrophils during GAS infection was measured using the LEGENDplex™ human inflammation cytometric bead assay over 360 min. Neutrophils differentially released (C) IL-1β, (D) TNF-α, (E) IL-8 and (F) IL-18 in response to GAS infection (duplicate measurements, n=6 donors). (B-F) Results are the pooled means±SD. *p<0.05, **p<0.01, ***p<0.001 and ****p<0.0001.

The above data suggest both GAS strains induce pyroptosis in neutrophils, but the response to 5448AP is delayed. To explore this possibility further, human neutrophils were incubated with 5448, 5448AP or in the absence of GAS, and caspase-1 activity assessed using a fluorogenic flow cytometric caspase-1 assay (Figures 3B and S3A-C). 5448 infected neutrophils showed increased active caspase-1 at 60 min and 180 min (p<0.05), whilst 5448AP infection increased caspase-1 at 180 min (p<0.05) compared to uninfected neutrophils (Figure 3B). No significant differences were observed between strains although caspase-1 activity was slightly reduced during 5448AP infection compared to 5448 (Figure 3B), similar to the immunoblot data for caspase-1 p46 (Figures 3A and S2C). Thus, GAS infection of human neutrophils corresponds to increased caspase-1 activity and caspase-4 expression, consistent with inflammasome activation and pyroptosis.

### Proinflammatory cytokines IL-1β and TNF-α are released by neutrophils in response to GAS infection

The release of the inflammatory cytokine IL-1β occurs due to caspase-1 activation and is associated with the induction of pyroptosis (Broz et al., 2010). Uncoordinated release of various inflammatory cytokines can exacerbate infection (Tecchio et al., 2014). During GAS infection, disease severity is negatively correlated to cytokine concentration (Norrby-Teglund et al., 2000). Therefore, human neutrophils were incubated with 5448, 5448AP or in the absence of GAS and supernatants assessed for cytokines using a flow cytometric bead assay. Neutrophils incubated in the absence of GAS failed to release IL-1β (Figure 3C) and TNF-α (Figure 3D) despite releasing increasing amounts of IL-8 over 360 min (Figure 3E). Incubation with GAS induced IL-1β release from neutrophils at 180 min and 360 min, with significantly greater release during 5448AP infection compared to 5448 at 180 and 360 min (p<0.05, p<0.0001, Figure 3C). Incubation with GAS also induced significant release of TNF-α from neutrophils at 180 min (p<0.001) and 360 min (p<0.0001), with significantly greater release during 5448 infection compared to 5448AP at 360 min (p<0.0001, Figure 3D). Compared to uninfected neutrophils, 5448 induced significant release of IL-8 from neutrophils at 60 min and 180 min (p<0.05), with significantly greater release than 5448AP at 60 min (p<0.05, Figure 3E). Neutrophils released minimal amounts of IL-18 over 360 min, and this response did not change following infection with both 5448 or 5448AP (Figure 3F), suggesting that neutrophils have no major role in IL-18 release over 360 min *in vitro*. These data indicate that both 5448 and 5448AP induce IL-1β release from human neutrophils, consistent with the hypothesis that both GAS strains induce pyroptosis in these cells. Moreover, the results indicate differential cytokine release from human neutrophils infected with GAS, with 5448AP inducing greater IL-1β release and 5448 inducing greater TNF-α and IL-8 release.

### GAS infection may cause neutrophil dysfunction

Neutrophil adhesion and activation are central to the innate immune response, with CD11b (Mac- and CD66b (CEACAM8) playing key respective roles (Arnaout, 1990, Power et al., 2001, Skubitz et al., 1996). CD16 (FcγRIII) facilitates opsonisation of pathogens and is down-regulated during apoptosis (Dransfield et al., 1994, Fossati et al., 2002), whilst down-regulation of CD31 (PECAM-1) facilitates clearance of apoptotic neutrophils and the resolution of inflammation (Brown et al., 2002, Kurosaka et al., 2003). Therefore, to further explore the impact of GAS on neutrophil function, human neutrophils were incubated with 5448, 5448AP or in the absence of GAS and the expression of cell-surface CD11b, CD66b, CD16 and CD31 was assessed using flow cytometry (Figure 4A-F). In the absence of GAS, neutrophils displayed minor increases in CD11b expression over time, while incubation with GAS resulted in minor decreases over time, with 5448 inducing a greater loss of CD11b than 5448AP, though these differences were not statistically significant (p=0.289, Figure 4C). Neutrophils incubated in the absence of GAS revealed minor increases in cell-surface CD66b expression over time (Figure 4D). Incubation with GAS further increased expression of CD66b, with 5448 inducing a slightly greater increase than 5448AP, though these differences were not statistically significant (p=0.162, Figure 4D). Incubation of neutrophils in the absence of GAS resulted in a steady decline in CD16 expression over time, a loss significantly increased by co-incubation with either GAS strain 180 min post-infection (p<0.0001, Figure 4E). Notably, incubation of neutrophils with 5448 induced a significantly greater loss of CD16 expression than 5448AP incubation at all earlier time points (Figure 4E). Similar to CD16, incubation of neutrophils in the absence of GAS resulted in a steady decline in CD31 expression over time, a loss significantly increased by co-incubation with either GAS strain 180 min post-infection (p<0.0001, Figure 4F). Moreover, incubation of neutrophils with 5448 induced a significantly greater loss of CD31 expression than 5448AP incubation at earlier time points. Collectively, these data indicate that GAS has minimal impact upon human neutrophil CD11b and CD66b, but increases the loss of CD16 and CD31 expression, which may impact opsonisation of GAS and subsequent efferocytosis of neutrophils by macrophages, during infection.

**Figure 4.**
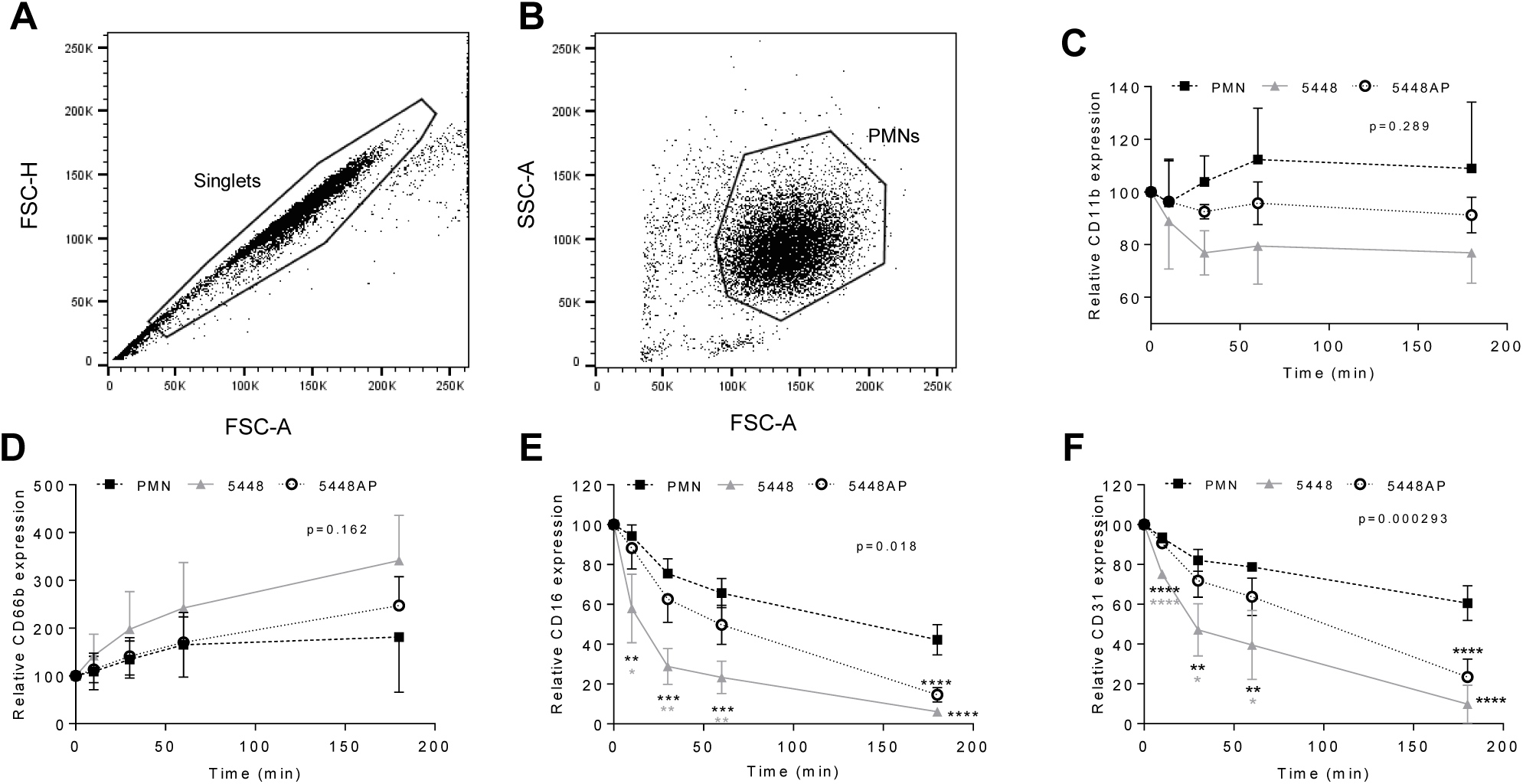
GAS infection changes neutrophil functionality. Human neutrophils (PMNs) were analysed using flow cytometry, sequentially gated upon (A) ‘Singlets’ (FSC-A vs. FSC-H, 10,000 events collected) then (B) viable ‘PMNs’ (FSC-A vs. FSC-H). Purified neutrophils infected with GAS were sampled over 180 min for the cell-surface expression of (C) CD11b-FITC (n=4 donors), (D) CD66b-PerCP/Cy5.5 (n=5 donors), (E) CD16-FITC (n=4 donors) and (F) CD31-PE/Cy7 (n=4 donors). Results are the pooled means±SD. *p<0.05, **p<0.01, ***p<0.001 and ****p<0.0001, with black asterisks denoting significance from control and grey asterisks between 5448 and 5448AP.

### GAS infection activates caspase-1 in neutrophils *in vivo*

The above data indicate that GAS influences human neutrophils *in vitro*. To explore if GAS may alter neutrophils *in vivo*, we assessed the neutrophil response in a mouse model of GAS infection (Ly et al., 2014, Maamary et al., 2010, Tsatsaronis et al., 2014). C57BL/6J mice were injected intradermally with 5448, 5448AP or saline and euthanised at 6 or 24 hours post infection, with neutrophil populations (CD45^+^/CD11b^+^/Ly-6G^+^), caspase-1 activation and CD16 expression characterised by flow cytometry (Figures 5A-C, 5E and S4A-F). Infection of mice with either GAS strain significantly increased neutrophil populations at sites of injection at both 6 and 24 h compared to saline control (p<0.001, Figure 5A). The percentage of circulating neutrophils in the blood at 6 h significantly increased during infection with both GAS strains (p<0.01), though the response to 5448AP was significantly greater than 5448 (p<0.05, Figure 5B). Blood neutrophil populations returned to baseline after 24 h, indicating the importance of neutrophils during the early stages of infection (Figure 5B). Murine neutrophils lavaged from sites of saline injection had low amounts of active caspase-1 (Figure 5B). Infection with 5448AP, but not 5448, significantly increased caspase-1 activation at 6 h (p<0.05), while both strains increased caspase-1 activation at 24 h (p<0.001, Figure 5C). Cytokines were assessed in murine serum using a flow cytometric bead assay. IL-1β concentrations remained low for saline control and 5448 infected mice but were significantly increased at 24 h during 5448AP infection (p<0.0001, Figure 5D). Neutrophils lavaged from the site of saline and GAS injection displayed CD16 expression, with CD16 expression significantly increased due to 5448 but not 5448AP infection at 6 h when compared to saline control (p<0.05, Figure 5E). The expression of CD16 on neutrophils from mice infected with 5448 significantly decreased between 6 and 24 h (p<0.01), but no significant differences were noted between times for either saline or 5448AP groups (Figure 5E).

**Figure 5.**
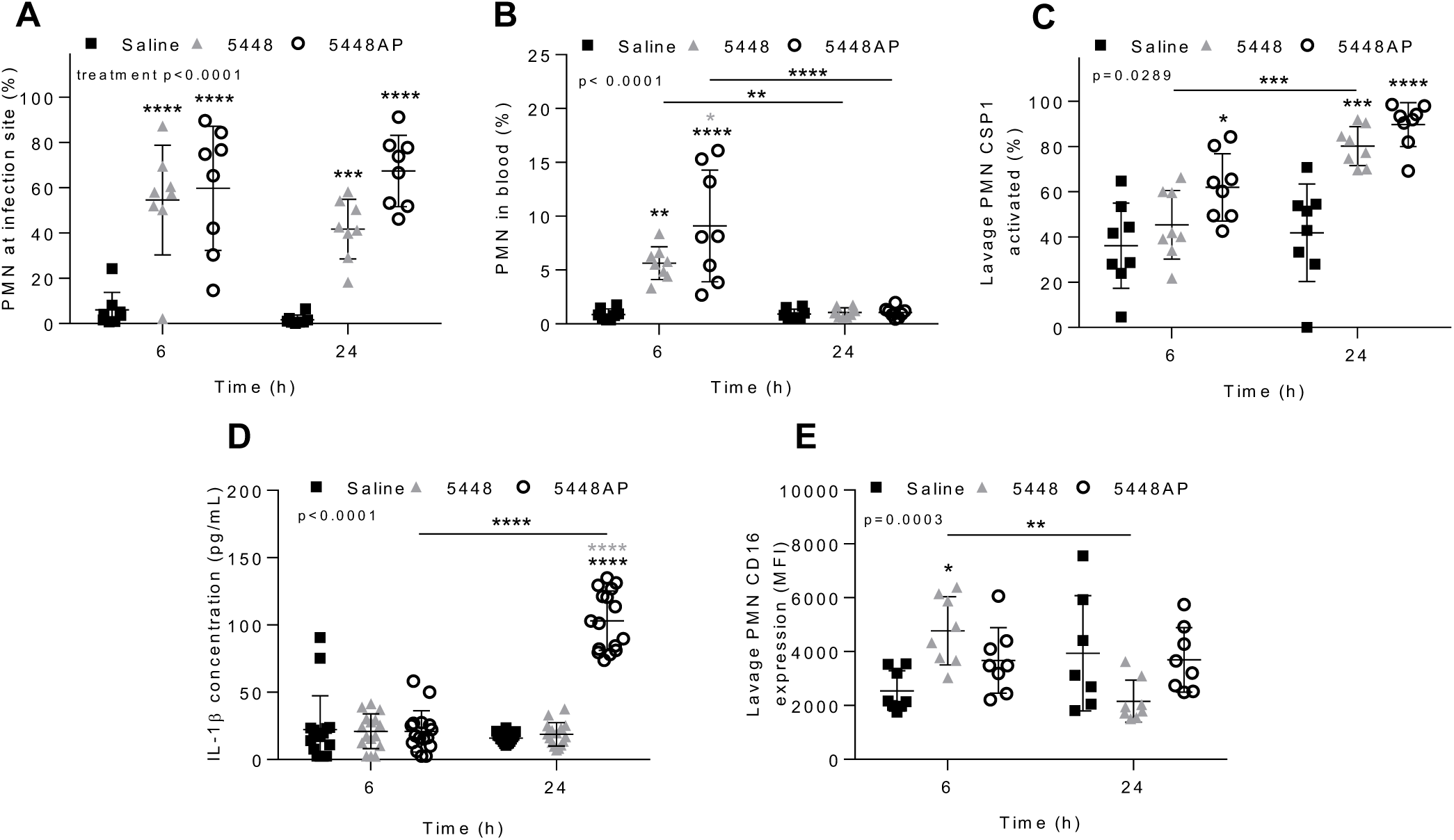
GAS infection *in vivo* activates caspase-1 in murine neutrophils. Percentage of CD45^+^ cells (A) lavaged from site of injection and in (B) circulating blood characterised as neutrophils (PMNs, CD11b^+^/Ly-6G^+^) by flow cytometry (Figure S4A-E). (C) Caspase-1 activation in neutrophils at site of GAS infection was confirmed via flow cytometry using FLICA 660 (660-YVAD-FMK, Figure S4F). Each data point represents the results from a single mouse where n=8. (D) The release of IL-1β in mouse serum during GAS infection was measured using the LEGENDplex™ mouse inflammation cytometric bead assay at 6 and 24 h post infection where duplicate measurements are shown and n=8. (E) CD16 expression of PMNs lavaged from the site of injection and determined using flow cytometry. Each data point represents the results from a single mouse where n=8. Results are the means±SD. *p<0.05, **p<0.01, ***p<0.001 and ****p<0.0001, with black asterisks denoting significance from control and grey asterisks between 5448 and 5448AP, or as indicated by a line.

## Discussion

Here we have mapped the human neutrophil response to GAS, addressing cell-death, signalling and the inflammatory profile during the early stages of M1T1 (5448 and 5448AP) infection, while also confirming *in vitro* findings in a murine intradermal GAS infection model. For the first time, we describe an inflammatory neutrophil phenotype, as evidenced by caspase-1 activation *in vitro* and *in vivo*, that may promote inflammation and exacerbate GAS disease. Most notably, changes to the expression of cell-surface proteins CD16 and CD31 may impede the removal of bacteria and clearance of neutrophils at sites of infection, hindering the resolution of inflammation. Further, we elucidate differences between 5448 and 5448AP that aid in understanding the survival and proliferation of GAS harbouring *covS* mutations in the presence of human neutrophils.

The temporal expression of neutrophil cell surface markers has not previously been investigated in the context of GAS infection. Our data show that significant reductions to both CD16 and CD31 occur in neutrophils due to GAS infection *in vitro*. Additionally, *in vivo* data indicates that a reduction in CD16 expression is seen during 5448 infection between 6 and 24 h. CD16 functions as a receptor for opsonised bacteria and immune complexes (Fossati et al., 2002), whilst the loss of CD16 has also been associated with the induction of apoptosis (Dransfield et al., 1994). The induction of apoptosis following bacterial phagocytosis is advantageous to the host as it promotes controlled removal of bacteria and the resolution of inflammation (Kobayashi et al., 2018). We hypothesise that the retention of CD16 during infection promotes a reduction in apoptosis and prolonged degranulation at the site of infection. Further, CD31 expression is reduced during GAS infection, facilitating the eventual removal of neutrophils from sites of infection (Kobayashi et al., 2003). Parallel to CD16, CD31 is retained during 5448AP infection of neutrophils to a greater extent than 5448 infection *in vitro*. This may result in neutrophils remaining at sites of infection longer, further increasing inflammation. Additionally, we report a trend towards increased cell-surface expression of CD11b and CD66b during GAS infection. Migration of neutrophils to sites of infection, facilitated by CD11b-mediated neutrophil adhesion (Arnaout, 1990), may therefore be affected during GAS infection. Down-regulation of CD11b can prevent accumulation of neutrophils at inflammatory sites (Huston et al., 2009), however retention, such as that seen during 5448AP infection may contribute to prolonged inflammation. Neutrophil activation can be determined through the expression of cell-surface CD66b following stimulation (Skubitz et al., 1996). Reduced cell-surface CD66b expression in response to 5448AP infection may therefore indicate a reduction to neutrophil function and further explain the resistance to neutrophil-mediated killing. Collectively, the neutrophil response is altered and inflammation prolonged at sites of *covS* mutant GAS infection.

Disruption of the role neutrophils play during the resolution of GAS infection and induction of inflammatory death pathways has been reported previously (Kobayashi et al., 2003, Tsatsaronis et al., 2015). Proinflammatory activation of the NLRP3 inflammasome in macrophages has been demonstrated for M1 GAS (Valderrama et al., 2017), here we provide the first evidence that GAS activates caspase-1 in human and murine neutrophils, consistent with inflammasome activation and pyroptosis. Caspase-1 activation was consistent during infection with both M1T1 GAS isolates, however the release of inflammatory cytokine IL-1β, a hallmark of clinical invasive infection (LaRock and Nizet, 2015), was only significantly increased in response to 5448AP infection *in vitro* and *in vivo*. In murine bone marrow-derived dendritic cells caspase-1 activation is essential for IL-1β release (Schneider et al., 2017). The activation of caspase-1 in human and murine neutrophils does not correlate directly to the amount of IL-1β released, and an alternative mechanism may facilitate IL-1β maturation. Type I interferons are known regulators of IL-1β in GAS infected mice (Castiglia et al., 2016), where increased IL-1β release in murine serum seen in this study may also be attributed to disruption of this homeostatic relationship. Increased IL-1β, being a neutrophil chemoattractant, therefore may further promote inflammation at sites of infection following release (Chen et al., 2007), supporting the hypothesis that *covS* mutant GAS promote greater levels of inflammation during GAS infection.

Previous studies have characterised differences between 5448 and 5448AP, demonstrating resistance to neutrophil-mediated killing *in vitro* and increased bacterial dissemination and decreased survival *in vivo* following the acquisition of *covS* mutations (Fiebig et al., 2015, Maamary et al., 2010, Walker et al., 2007). Here, we demonstrate that 5448AP is not only resistant to neutrophil-mediated killing but proliferates in the presence of functional neutrophils. GAS survival has been demonstrated intracellularly in murine neutrophils and human macrophages (Medina et al., 2003a, Medina et al., 2003b, O’Neill et al., 2016) and can even be facilitated by host red blood cells (Wierzbicki et al., 2019). Increased 5448AP proliferation may result from the reduction in neutrophil ROS response during infection. GAS has previously been shown to display resistance to ROS through numerous virulence mechanisms (Henningham et al., 2015). The absence of a hostile environment may further permit bacterial replication. Additionally, the increased production of ROS can induce neutrophil death via apoptosis (Geering and Simon, 2011). Reduced neutrophil ROS production during 5448AP infection may therefore explain the reduction seen in caspase-3 activation (apoptosis). Previously it has been demonstrated that GAS M1 strain MGAS5005 modulates neutrophil apoptosis (Kobayashi et al., 2003), a finding which is supported by the current study.

Changes to the function of neutrophils during the initial innate immune response have the potential to hinder mechanisms essential for the resolution of infection. Disruption to these processes during GAS infection may induce this neutrophil phenotype we have described, with increased inflammatory properties and potential to exacerbate infection. Caspase-1 activation during GAS infection of neutrophils indicates possible inflammasome activation and death via pyroptosis. Further studies should explore the precise cell death mechanism. A reduction in apoptosis is evident during *covS* GAS infection, as has been reported for non-M1T1 GAS (Tsatsaronis et al., 2015), whilst modulation of apoptosis has previously been implicated as a factor contributing to GAS survival (Kobayashi et al., 2003). The current study aids in understanding the complex host-pathogen interaction identifying host factors that may contribute to severe pathology of invasive GAS infection. Future studies may be able to isolate host-specific targets that could be exploited to control inflammation and tissue damage during invasive GAS infection.

## Acknowledgements

We thank Professor Marijka Batterham (University of Wollongong) for her statistical consultation. We thank Associate Professor Thomas Proft (University of Auckland, New Zealand) for pLZ12Km2-P23R:TA. We also thank our generous blood donors for their participation and time.

## Author Contributions

Conceptualisation, M.S.S, R.S., J.A.T., J.G.W.; Methodology, M.S.S., R.S., J.G.W., J.G., J.D.M., D.L.; Investigation, J.G.W., D.L., N.J.G., H.K.N.V.; Data and figure curation, J.G.W., D.L., R.S., M.S.S.; Writing-Original draft, J.G.W.; Writing-Review and editing, M.S.S., R.S., J.G.W., D.L., N.J.G., J.D.M., H.K.N.V., J.G., J.A.T.; Funding acquisition, M.S.S and D.L.; Resources, M.S.S., J.D.M.; Supervision, M.S.S., R.S., J.G.

## Declaration of Interests

The authors declare no competing interests.

## Materials and Methods

### Ethics statement

All experiments involving the use of human blood were conducted with informed consent of healthy volunteers, approved and authorised by the University of Wollongong Human Research Ethics Committee (Protocol HE08/250). All experiments involving the use of animals were approved and authorised by the University of Wollongong Animal Ethics Committee (Protocol AE18/10).

### Bacterial strains and culture

*Escherichia coli* MC106 were grown in Luria–Bertani Broth (LB) at 37 °C with constant shaking. M1T1 *S. pyogenes* invasive clinical isolate 5448 (*emm1*) and hypervirulent animal passaged 5448AP (containing a non-functional control of virulence regulator mutation) have been described previously (Aziz et al., 2004, Chatellier et al., 2000). GAS strains were routinely cultured at 37°C on horse-blood agar (Thermo Fisher) and enumerated on yeast supplemented (1% w/v) Todd-Hewitt (THY, Benton Dickson) agar. Static overnight cultures were grown at 37°C in THY then sub-inoculated (1:10) into fresh THY and grown to mid-logarithmic phase, antibiotics were added when required (100 μg/ml ampicillin and 200 μg/ml kanamycin). Bacterial pellets were washed twice with phosphate buffered saline (PBS) then resuspended at the specified multiplicity of infection (MOI) and assay specific media conditions.

### Construction of GFP expression vector pLZ12Km2-P23R-TA:GFP for GAS

In brief, stable GFP expression by GAS was created by synthesizing the ribosomal binding site (RBS) and *gfp* gene from pDC*erm*-GFP (Ly et al., 2014) into the pUC57 plasmid (GenScript), resulting in pUC57-RBSGFP plasmid. RBS and *gfp* from pUC57-RBSGFP were then sub-cloned using NotI into the toxin–antitoxin (TA) stabilized expression plasmid pLZ12Km2-P23R:TA, kindly provided by Associate Professor Thomas Proft (University of Auckland, New Zealand) (Loh and Proft, 2013), to produce pLZ12Km2-P23R-TA:GFP. The resultant plasmid system utilises the *Streptococcal* ω–ε–ζ TA cassette to achieve segregational plasmid stability under non-selective conditions (Lioy et al., 2010).

Specifically, plasmids pLZ12Km2-P23R:TA and pUC57-RBSGFP (engineered with RBS and *gfp* from pDC*erm*-GFP) were first transformed into chemically competent *E. coli* MC106 via electroporation. The plasmids were then retrieved and purified from *E. coli* cultures using a Wizard® Plus SV Minipreps DNA Purification System (Promega). The pLZ12Km2-P23R:TA plasmid was digested with 20 U NotI enzyme and further treated with 5 U shrimp alkaline phosphatase. The pUC57-RBSGFP plasmid was also incubated with 20 U NotI to excise the RBS and *gfp* gene from pUC57-RBSGFP, and these were then ligated into digested pLZ12km2-P23R:TA plasmid. The resulting plasmid, pLZ12Km2-P23R-TA:GFP, was then transformed and cloned in MC106 *E. coli*, and later extracted and purified. pLZ12Km2-P23R-TA:GFP was transformed into GAS using standard GAS electroporation techniques (McLaughlin and Ferretti, 1995). Transformed GAS was confirmed for GFP expression via flow cytometry.

### Isolation of human neutrophils

Venous human blood was drawn into 10 mL lithium heparin-coated Vacutainer (Benton Dickson) tubes and layered over equal volumes of Polymorphprep (Axis Shield) and centrifuged as per manufacturer’s instructions. The resulting layer of neutrophils was isolated and erythrocytes hypotonically lysed. Isotonic concentration was restored with Hank’s Balanced Salt solution (without Ca^2+^ or Mg^2+^, Corning Inc.). Prior to experimentation neutrophils were resuspended at specified concentrations in complete medium, Roswell Park Memorial Institute (RPMI)-1640 medium (Life Technologies) containing 2% heat-inactivated autologous plasma and 2 mM L-glutamine (Life Technologies), unless otherwise stated. Neutrophil viability was assessed via Trypan Blue (Sigma-Aldrich) staining and neutrophil purity was assessed using a Benton Dickson LSR Fortessa X-20 flow cytometer via distinct forward and side scatter profiles or CD66b-peridinin chlorophyll protein (PerCP)/Cy5.5 (clone G10F5, BioLegend) expression. Neutrophils were maintained at room temperature throughout processing.

### *In vitro* infection of neutrophils with GAS

Purified human neutrophils were seeded in either 96-well plates at MOI (GAS:neutrophil) 1:10 (survival and proliferation) or 10:1 (ROS production and cytokine release), 24-well plates at 10:1 (phagocytosis, Annexin-V/Viability staining, CD expression and FAM-FLICA caspase-1 activation) or 6-well plates at 10:1 (immunoblotting) and incubated at 37°C in 5% CO_2_.

### GAS survival and proliferation

Human neutrophils were co-cultured with GAS or lysed via three freeze-thaw cycles prior to incubation. To block phagocytosis neutrophils were preincubated with 10 μM cytochalasin D (Cayman Chemicals) in complete medium at 37°C for 30 min prior to infection. At indicated time points neutrophils were hypotonically lysed and surviving bacteria serially diluted before plating on THY agar.

### GAS phagocytosis

Human neutrophils were co-cultured with GAS expressing GFP for the times indicated in complete medium. Cells were removed and washed twice with 10% (v/v) heat-inactivated foetal bovine serum (FBS, Bovogen Biologicals) diluted in PBS, followed by data acquisition via flow cytometry.

### ROS production

ROS production was assessed as previously described (Kobayashi et al., 2003). Briefly, prior to infection, human neutrophils were incubated in complete medium containing 25 μM dichlorofluorescein (DCF, Molecular Probes) for 45 min at room temperature (RT). ROS production was measured fluorometrically (ex485 nm em520 nm) using a POLARstar Omega plate reader (BMG Labtech).

### Flow cytometry

For human neutrophil in vitro assays, flow cytometry data was acquired using a BD LSR Fortessa X-20 with excitation lasers; violet (405 nm), blue (488 nm), yellow/green (561 nm) and red (640 nm). Bandpass filters for Zombie Aqua Fixable Viability Kit (525/50), fluorescein isothiocynate (FITC)/GFP/FAM FLICA (525/50), PerCP-Cy5.5 (695/40) R-phycoerythrin (PE, 586/15), propidium iodide (PI, 610/20), PE-Cy7 (780/60) and allophycocyanin (APC, 670/30) were used. For murine infection studies, flow cytometry data was acquired using an Invitrogen Attune NxT with excitation lasers; violet (405 nm), blue (488 nm), yellow (561 nm) and red (638 nm). Bandpass filters VL1 (440/50), VL2 (512/25), BL1 (530/30), YL1 (585/16), YL3 (695/40) and RL2 (720/30) were used. Data was analysed using FlowJo software V10.6.1 (TreeStar Inc.).

### Annexin-V/viability staining

To assess cell viability human neutrophils at indicated times were washed with PBS and incubated with Zombie Aqua Fixable Viability Kit (BioLegend) for 15 min at RT. Cells were washed again once with PBS, then 10% (v/v) FBS in PBS and stained with Annexin-V-FTIC (BioLegend) for 15 min at RT. Cells were analysed immediately via flow cytometry.

### Immunoblotting and antibodies

Following co-culture in the presence or absence of GAS, human neutrophil lysates were prepared in RIPA buffer (150 mM NaCl, 5 mM EDTA, 50 mM Tris, 1.0% (v/v) Triton X-100, 0.1% (w/v) SDS, 0.5% (w/v) sodium deoxycholate, 2x cOmplete protease inhibitor cocktail (Roche), 1 mM phenylmethylsulphonyl fluoride, 5 mM sodium pyrophosphate, 5 mM sodium molybdate and 5 mM β-glycerophosphate) following co-culture in the presence or absence of GAS and incubated for 30 min on ice with intermittent vortexing. Soluble fractions were separated via centrifugation at 4°C, snap frozen in liquid N_2_ and stored at −80°C. Protein concentrations were determined by comparison to bovine serum albumin standards using the DC Protein Assay (Bio-Rad) and absorbance (A^750nm^) measured using a POLARstar Omega plate reader. Neutrophil lysates (20 μg) were separated on 4-20% TGX Stain-Free protein gels (Bio-Rad) as per the manufacturer’s running conditions then activated for 5 min under UV to determine total protein (Bio-Rad ChemiDoc XR, Image Lab Software). Proteins were transferred to PVDF membranes (Bio-Rad), blocked, then incubated overnight at 4°C with caspase-1 polyclonal (1:1000, #2225), caspase-3 polyclonal (1:1000, #9662), caspase-4 polyclonal (1:1000, #4450) or caspase-8 monoclonal (1:1000, #4790) antibodies (Cell Signalling Technology). PVDF membranes were washed thrice between incubations for 5 min with tris-buffered saline containing 0.1% (v/v) Tween 20. PVDF membranes were incubated with horseradish peroxidase-conjugated goatαrabbit IgG (1:5000, Invitrogen) for 1 h at RT and detected using Clarity or Clarity Max Western ECL Blotting Substrate (Bio-Rad) and imaged (Amersham AI600). Bands were quantified using ImageJ software (National Institutes of Health) as area under the peak and normalised over UV determined total protein.

### FAM-FLICA caspase-1 activation

Caspase-1 activation was measured using FAM-FLICA Caspase-1 assay kit (ImmunoChemistry Technologies) as per the manufacturer’s instructions. In brief, following co-culture of human neutrophils with GAS, cells were removed and incubated in serum free medium (RPMI-1640) containing 1 x FAM-YVAD-FMK for 60 min at 37°C. Cells were then analysed for caspase-1 activation via flow cytometry.

### Cytometric cytokine bead assay

Human neutrophil supernatants were at indicated times during GAS infection, snap frozen with liquid N_2_ and stored at −80°C for a period no longer than 5 days. Neutrophils were assessed for the release of cytokines IL-1β, IL-8, IL-18 and TNF-α using the LEGENDplex™ human inflammation panel bead-based immunoassay (BioLegend) and flow cytometry as per manufacturer’s instructions. In brief, samples and standards were incubated with beads and detection antibodies for 120 min at room temperature, with shaking (600 rpm) protected from light. PE-conjugated streptavidin was then added and incubated for a further 30 min at room temperature, with shaking (600 rpm) protected from light. Beads were washed twice with LEGENDplex™ wash buffer and data collected via flow cytometry. Beads were initially gated upon FSC-A and SSC-A (A and B populations). Cytokines have signature APC fluorescence with quantitative expression determined using PE fluorescence in comparison to cytokine standards. Data was analysed using the LEGENDplex™ software V8.0 (VigeneTech Inc.).

### Cluster of differentiation expression

Human neutrophil cell surface CD expression was assessed during GAS infection. Neutrophils at indicated times were washed with PBS then routinely incubated with Zombie Aqua Fixable Viability Kit for 15 min at RT. Cells were washed once again in PBS, followed by 10% (v/v) FBS in PBS. Neutrophils were stained with fluorochrome-conjugated antibodies CD11b-FITC (Clone ICRF44), CD31-PE/Cy7 (Clone WM59), or CD66b-PerCP/Cy5.5 (Clone G10F5) (BioLegend) and CD16-FITC (Clone 3G8, Benton Dickson) for 15 min at RT. Neutrophils were washed with 10% (v/v) FBS in PBS then analysed via flow cytometry.

### Murine intradermal GAS infection model

Intradermal GAS challenge of C57BL/6J mice has been described previously (Ly et al., 2014, Maamary et al., 2010, Tsatsaronis et al., 2014). In brief, equal numbers of 6-8 week old male and female C57BL/6J mice (Australian BioResources) were anesthetised via isoflurane inhalation, followed by two intradermal injections into shaved left and right flanks with either 1 × 10^8^ CFU of mid-logarithmic GAS or 100 µL sterile 0.7% saline. Mice were euthanised via slow-fill CO_2_ asphyxiation at 6 or 24 h post-infection. Blood collected via cardiac puncture was separated into two aliquots for serum and flow cytometric analysis. Serum was collected by clotting for 1 h at RT then placed on ice until centrifugation at 1200 x *g* for 10 min at 4°C, then immediately stored at - 80°C until use. Remaining blood samples were mixed with 20 µL 0.5% sodium citrate (w/v) per mL of blood and placed on ice. Equal volume of PBS was added to the blood sample and centrifuged at 350 x *g* for 5 min. For erythrocyte lysis, samples were incubated with 1 mL of ammonium-chloride-potassium lysing buffer (150 mM NH_4_Cl, 1 mM KHCO_3_, 0.1 mM Na_2_CO_3_, pH 7.3) for 5 min with gentle agitation and centrifuged at 350 x *g* for 5 min. This process was repeated and cells were then washed with PBS and resuspended in 500 µL PBS and placed on ice. Additionally, sites of infection were lavaged with 1 mL of sterile 0.7% saline, and fluid collected from both right and left flanks was pooled to increase cell numbers. Pooled samples were kept cold until centrifugation at 350 x G for 5 min, resuspended in 500 µL of PBS and placed on ice. Both blood and lavage fluid samples were analysed via flow cytometry.

### Flow cytometric analysis of murine cells

Cell viability was assessed as described above. To assess caspase-1 activation and CD expression, cells were initially washed with 1 mL of PBS, followed by 1 mL of 10% (v/v) FBS in PBS. Cells were then centrifuged and simultaneously stained with 1 x 660-YVAD-FMK (FLICA 660 Caspase-1 assay kit, ImmunoChemistry Technologies) for caspase-1 activation and with fluorochrome-conjugated antibodies CD45-BV421 (clone 30-F11), CD11b-PE/Cy5 (clone M1-70) and Ly-6G-PE (clone 1A8) (Biolegend) and CD16-FITC (clone AT154-2, Bio-Rad) in RPMI 1640 media for 30 min at RT. Cells were washed with 10% (v/v) FBS in PBS and analysed via flow cytometry.

### Murine serum IL-1β concentration

IL-1β release was measured in murine serum using the LEGENDplex™ mouse inflammation panel bead-based immunoassay (BioLegend) and flow cytometry as described above except shaking was performed at 800 rpm.

### Statistical analyses

Graphs were created using Prism 6 (GraphPad Software Inc.) and statistical analysis was performed using Prism 6 and IBM SPSS Statistics 25 (IBM® corporation). Data was analysed using one-way and two-way ANOVA to determine significant differences and adjusted using Tukey HSD corrections or as stated otherwise. To account for the nested nature of the human neutrophil Annexin-V/Zombie and CD data with repeated measures over time in the same donor, a linear mixed model was used to determine significant interaction between treatment and time. Post-hoc tests were performed by multiple one-way ANOVA at each time point where multiple comparisons were adjusted using Tukey HSD corrections. *In vivo* data was analysed by two-way ANOVA where differences are shown between treatment and time, also using Tukey HSD corrections. Values presented for ‘p’ represent interaction (treatment*time) unless otherwise stated. * p<0.05; ** p<0.01; *** p<0.001; **** p<0.0001.

## Supplemental Information

**Supplementary figure 1.**
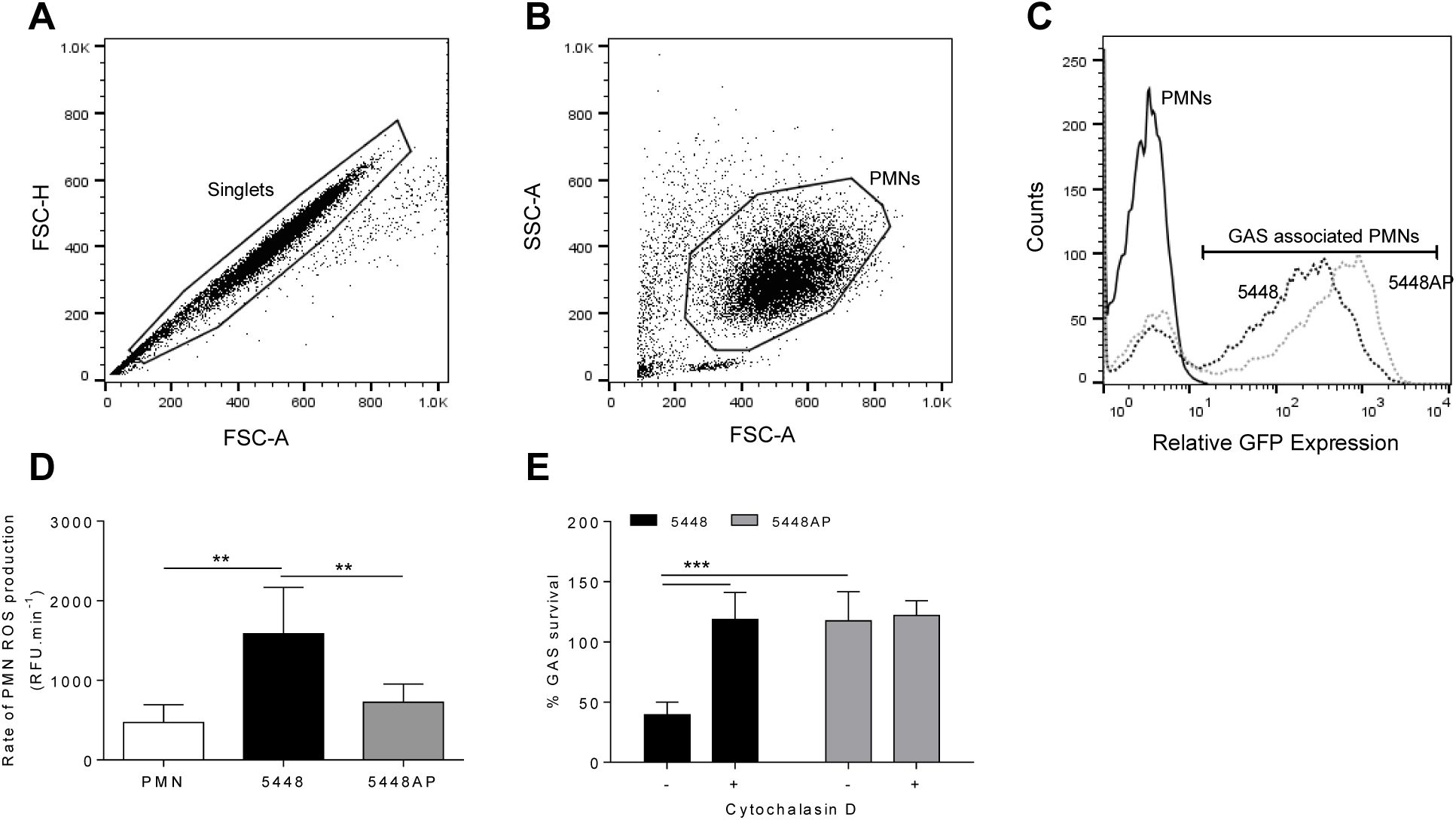
The association of fluorescent GAS (GFP) to human neutrophils determined via flow cytometry. (A) Primary neutrophil (PMN) ‘Singlets’ cell selection (FSC-A vs. FSC-H) and (B) viable ‘PMNs’ discrimination (FSC-A vs. SSC-A). (C) Representative histogram showing relative fluorescence (GFP) of GAS infected neutrophils and gating for the selection of GAS-neutrophil association. (D) Rate of ROS production by neutrophils between 30-60 min (n=6 donors). (E) GAS killing by human neutrophils is inhibited at 30 min following pre-incubation with actin polymerising molecule cytochalasin D (n=4 donors, Sidak’s multiple comparison). Results are the pooled means±SD (of triplicate measurements for panels D and E). **p<0.01, ***p<0.001.

**Supplementary figure 2.**
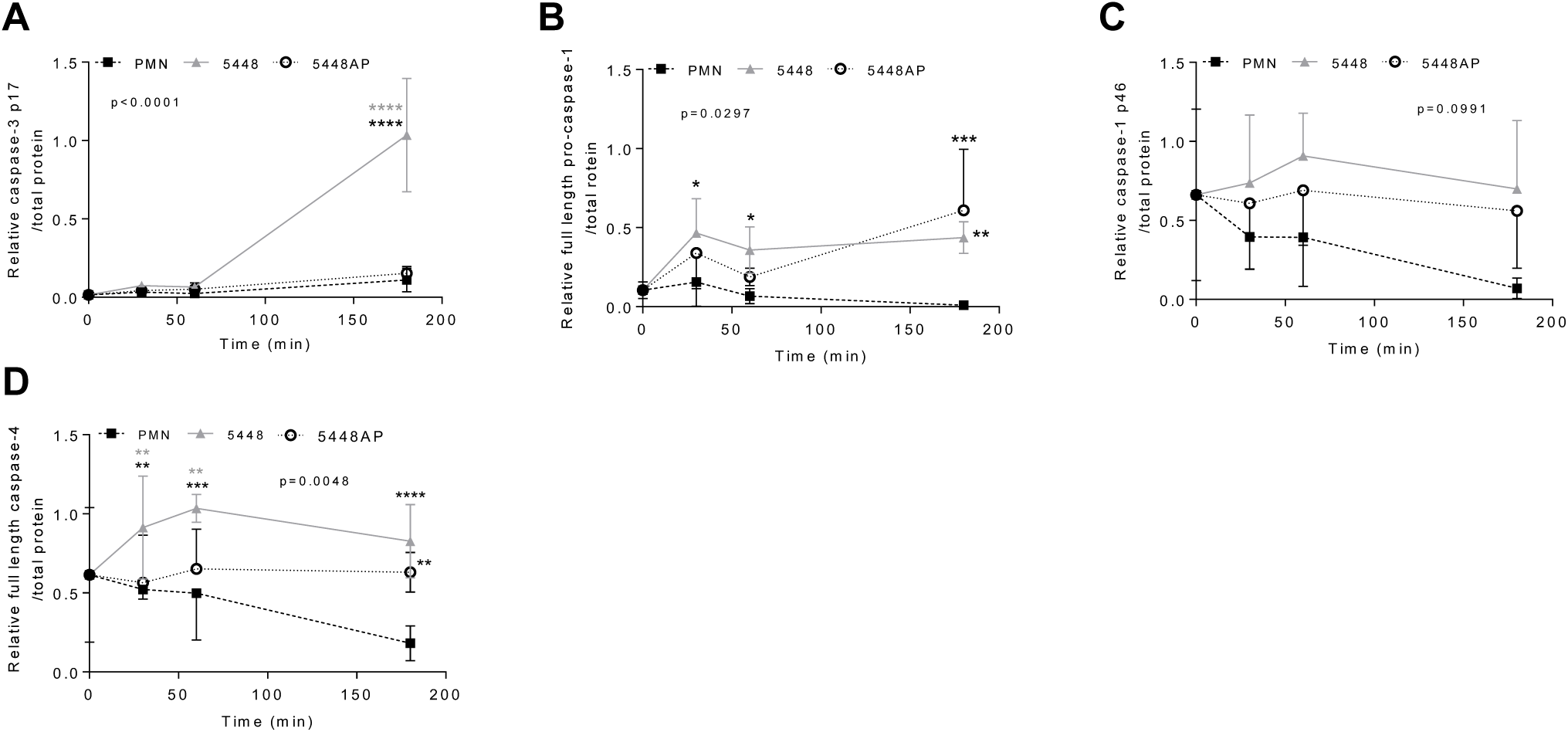
Quantification of protein bands in human neutrophil lysates identified through immunoblotting. (A) Caspase-3 p17, (B) caspase-1 full-length, (D) caspase-1 p46 and (D) caspase-4 immunoblots were imaged then analysed using ImageJ where area under the curve was normalised over total protein. Results are the means±SD where 3 donors were used. *p<0.05, **p<0.01, ***p<0.001 and ****p<0.0001, with black asterisks denoting significance to control and grey asterisks between 5448 and 5448AP.

**Supplementary figure 3.**
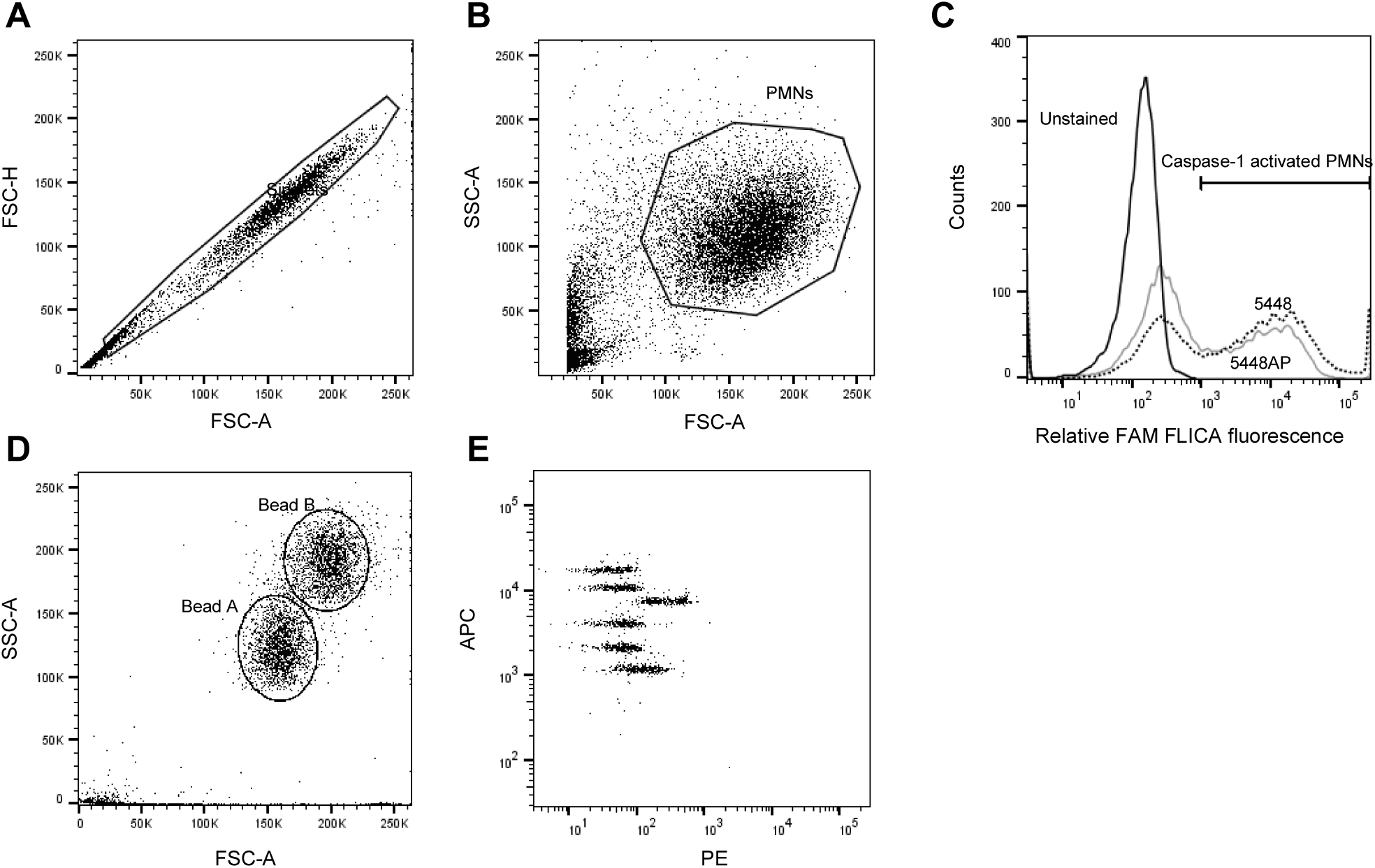
Human neutrophils were assessed for the activation of caspase-1 and release of cytokines via and flow cytometry. (A) Primary neutrophil (PMN) ‘Singlet’ cell selection (FSC-A vs. FSC-H) and (B) viable ‘PMNs’ discrimination (FSC-A vs. SSC-A). (C) Representative plot showing unstained and GAS infected neutrophils (Singlets/PMNs) with FLICA (FAM-YVAD-FMK) fluorescence. Gating strategy for LEGENDplex cytometric bead assay where (D) beads A and B were gated (FSC-A vs. SSC-A). Gates for beads A (shown) and B were assessed for (E) PE vs. APC fluorescence where cytokines had unique APC fluorescence and expression was measured via PE fluorescence when compared to a standard curve.

**Supplementary figure 4.**
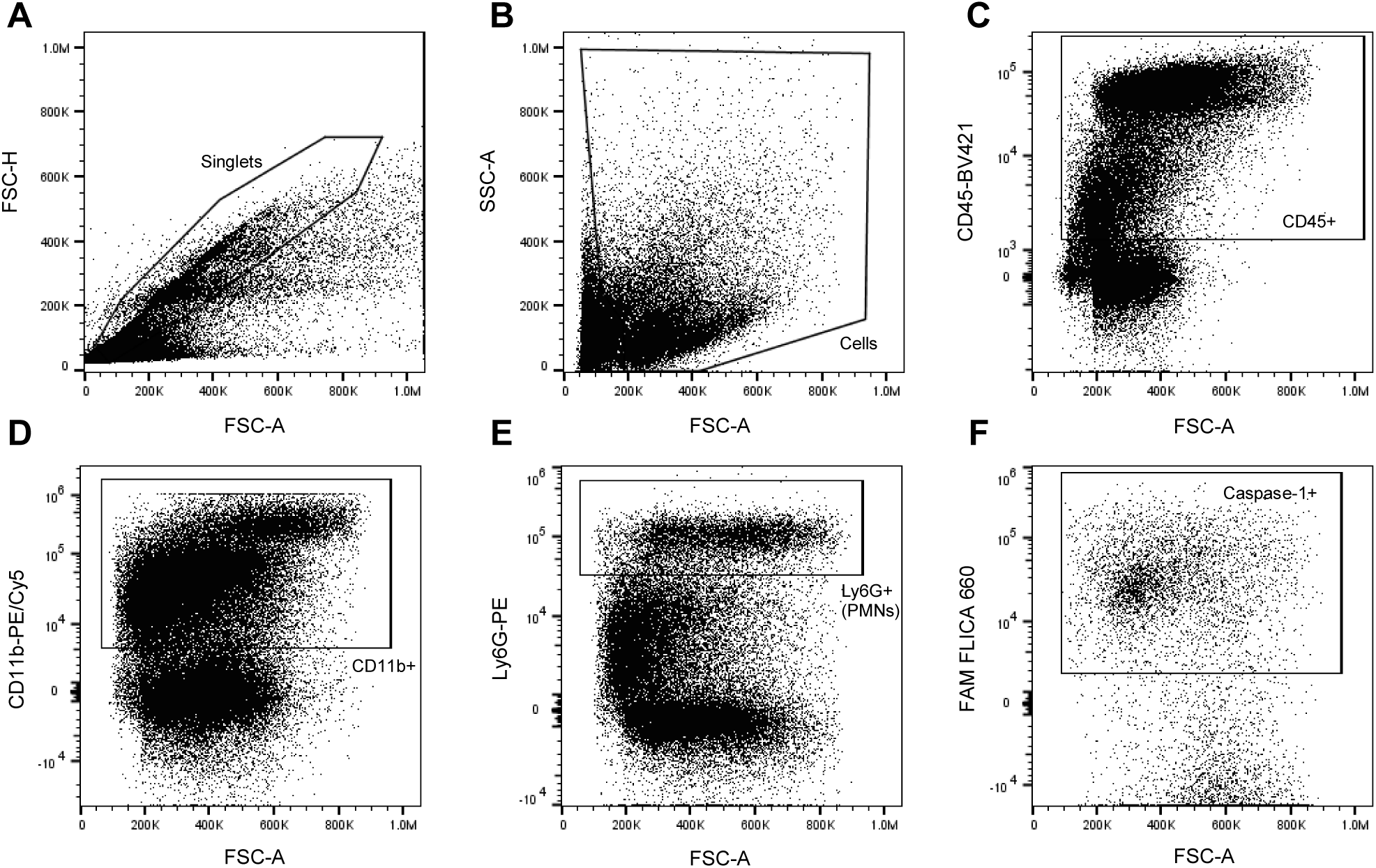
Flow cytometric sequential gating strategy for identification of murine neutrophils determining caspase-1 activation. (A) Primary ‘Singlets’ cell selection (FSC-A vs. FSC-H) and (B) ‘Cells’ discrimination (FSC-A vs. SSC-A) of murine blood and lavage fluid. ‘Cells’ were further gated upon (C) CD45-BV421 expression and (D) CD11b-PE/Cy5 expression. Neutrophils (PMNS) were defined as (E) Ly-6G-PE^+^ cells from the CD45^+^/CD11b^+^ population. Neutrophils were assessed for (F) caspase-1 activation using FLICA 660 (660-YVAD-FMK).

## Notes

### Competing Interest Statement

The authors have declared no competing interest.

### Summary of Updates

This revision of the manuscript has additional data about the functional role of neutrophils during GAS infection in vivo.

